# Instrumented individuals are not representative of the population. Survival of wolverines with and without GPS collars

**DOI:** 10.1101/2020.09.22.308007

**Authors:** Cyril Milleret, Richard Bischof, Pierre Dupont, Henrik Brøseth, John Odden, Jenny Mattisson

## Abstract

Results from biologging studies are often scaled up to population-level inferences and this begs the question: Are instrumented animals representative of the population given the potential bias in individual selectivity, the influence of capture, handling and wearing bio-loggers? The answer is elusive due to the challenges of obtaining comparable data from individuals without bio-loggers. Using non-invasive genetic data of a large carnivore, the wolverine (*Gulo gulo*) in Scandinavia, and an open-population spatial capture-recapture model, we were able to estimate and compare survival rates of individuals fitted with GPS collars versus individuals that had never been captured and collared. We found that mortality of GPS collared individuals was 47% lower compared to individuals without GPS collars. While instrumented and non-instrumented wolverines had similar risks of dying from legal culling, GPS collared individuals displayed lower mortality rates due to causes other than legal culling. Reports of positive effects of biologging on the fitness of individuals are rather uncommon and we argue that the most likely explanation is that GPS-collars could shield animals from poaching. Our results highlight the challenges of drawing population level inferences for large carnivores when using data from an instrumented sample of the population.

## Introduction

Telemetry and biologging systems have been instrumental in furthering our understanding of the ecology and cryptic behavior of wildlife. Technological development and miniaturization of devices have allowed researchers to track animals and obtain a wide range of previously unattainable information, such as fine scale movement data (Bischof et al. 2019), kill rates of predators (Gervasi et al. 2014), and habitat selection (Bouyer et al. 2015). Results from tracking studies are often scaled up to population-level inferences, with the assumption – explicit or implicit – that instrumented animals are representative of the population as a whole (Hebblewhite and Haydon 2010). There are two main reasons why this assumption may be violated: 1) the sample of instrumented individuals is non-random (i.e., a systematic bias in the selection of individuals studied) and 2) biologging and tracking themselves alter the biology of instrumented animals. Despite the long history and widespread use of wildlife telemetry applications, it is challenging to assess the reliability of telemetry studies for inferring processes and patterns in wild populations. This is largely due to the inherent catch-22: the methods whose impact is to be evaluated are often seen as the only reliable methods available for obtaining the parameters that are potentially impacted.

### Non-random sampling

Randomization is an essential requirement for inferring population properties from the properties of a sample. Yet, the selection of animal subjects enrolled in telemetry studies is rarely, if ever, fully randomized (Hebblewhite and Haydon 2010). For example, variation in the vulnerability to physical capture that is non-random with regards to biological attributes (such as behavior or fitness) is bound to lead to biased conclusions if estimates are extrapolated to the population level. Some deviations from representativeness may be easy to detect, such as a different age structure or gender ratio in the sample vs the population. Others, such as individual variation in boldness (Darrow and Shivik 2009) or social standing (Young et al. 2019) may not.

### Observer effect

Aside from animal welfare concerns (Putman 1995), methods that impact the study species will inadvertently introduce bias as the system studied is altered through the process of observation/data collection, the so-called “observer effect” (Canine 1990, MacFarlane and King 2002). First, capturing and handling are stressful for animals (Cattet et al. 2008, Esteruelas et al. 2016). Second, although benign in most cases (Bodey et al., 2018), attaching bio-logger devices can influence behavior or even have long-term detrimental effects on individuals (Lechenne et al. 2012, Bodey et al. 2018).

In addition to direct effects of capture, handling, and wearing biologging device, the fitness of instrumented individuals can be impacted indirectly if awareness of the presence of biologging devices alters human decision making in human-wildlife interactions. This may be particularly acute for rare, elusive, and controversial species such as large carnivores. Bio-logging is commonly used to study large carnivores and helps inform debate, policy, and management (Carricondo-Sanchez et al. 2020). Large carnivores are essential for ecosystem functioning but also get into conflict with humans as they prey on wild and domestic animals, and can occasionally injure or kill people (Packer et al. 2019). The resulting controversy and debate extend not only to large carnivore management, but also to the use of biologging in carnivore research. Indeed, this is not always perceived as positive by the public because captures are often costly and require the use of invasive techniques such as helicopter and anesthesia (Mech and Barber 2002). Furthermore, biologging can have indirect consequences for individuals as bio-loggers can interact with management decisions and lead to lethal actions that would have not occurred without the information provided by the logger (Meeuwig et al. 2015, Cooke et al. 2017). Real time information about individual location can also be made publicly available to avoid conflicts (Liberg et al. 2010) or be hacked by poachers (“cyber-poaching”,(Cooke et al. 2017)). Additionally, it has been suggested that instrumented animals could be protected from poaching due to the increased risk of detecting poaching events when animals wear a visible bio-logger. As a consequence, biologging of large carnivores may not only interfere directly with the behavior and fitness of individuals, but also indirectly by influencing management actions and illegal activities.

To investigate the effect of biologging, we used a decade worth of non-invasive genetic monitoring data of wolverines (*Gulo gulo*) in Scandinavia and compared survival rates of individuals with and without GPS collars using an open-population capture-recapture model. This unique data set contained data from individuals without capturing them, as both instrumented and non-instrumented individuals were sampled during monitoring. We show that instrumented large carnivores may not always constitute a representative sample of the population and call for prudence when extrapolating survival estimates from instrumented individuals to populations.

## Material and methods

### GPS collared individuals and study area

Wolverines were equipped with GPS collars after being darted from a helicopter and immobilized with medetomidine–ketamine, following pre-established protocols (Arnemo et al. 2012). GPS collars with GSM communication (GPS plus mini; Vectronic Aerospace GmbH, Berlin, Germany) were used. The collars were fitted with a release mechanism in the form of a cotton band that usually breaks apart after ∼0.5-2 years (depending on thickness of band and behavior of the animal). The handling protocols were approved by the Norwegian Experimental Animal Ethics Committee (FOTS ID 2826, FOTS ID 4699, FOTS 7017) and fulfilled their ethical requirements for research on wild animals. In addition, permits for wild animal capture were obtained from the Norwegian Environment Agency. Captures occurred in the winters (Jan-April) of 2010 to 2015. In total, we collared 43 individuals (♀21; ♂22), 18 in the Central (♀8; ♂10), and 25 in the Northern (♀13; ♂12) part of Norway (Appendix S1, Figure 1). Of these 18 (41%) where approximately 1-year old (average birthdate of wolverines 15^th^ of February). The wolverine population is distributed continuously between the two regions where individuals were captured (Bischof et al. 2019). However, we distinguished between the Central and Northern areas so that instrumented and non-instrumented individuals were exposed to similar ecological conditions.

**Figure 1.**
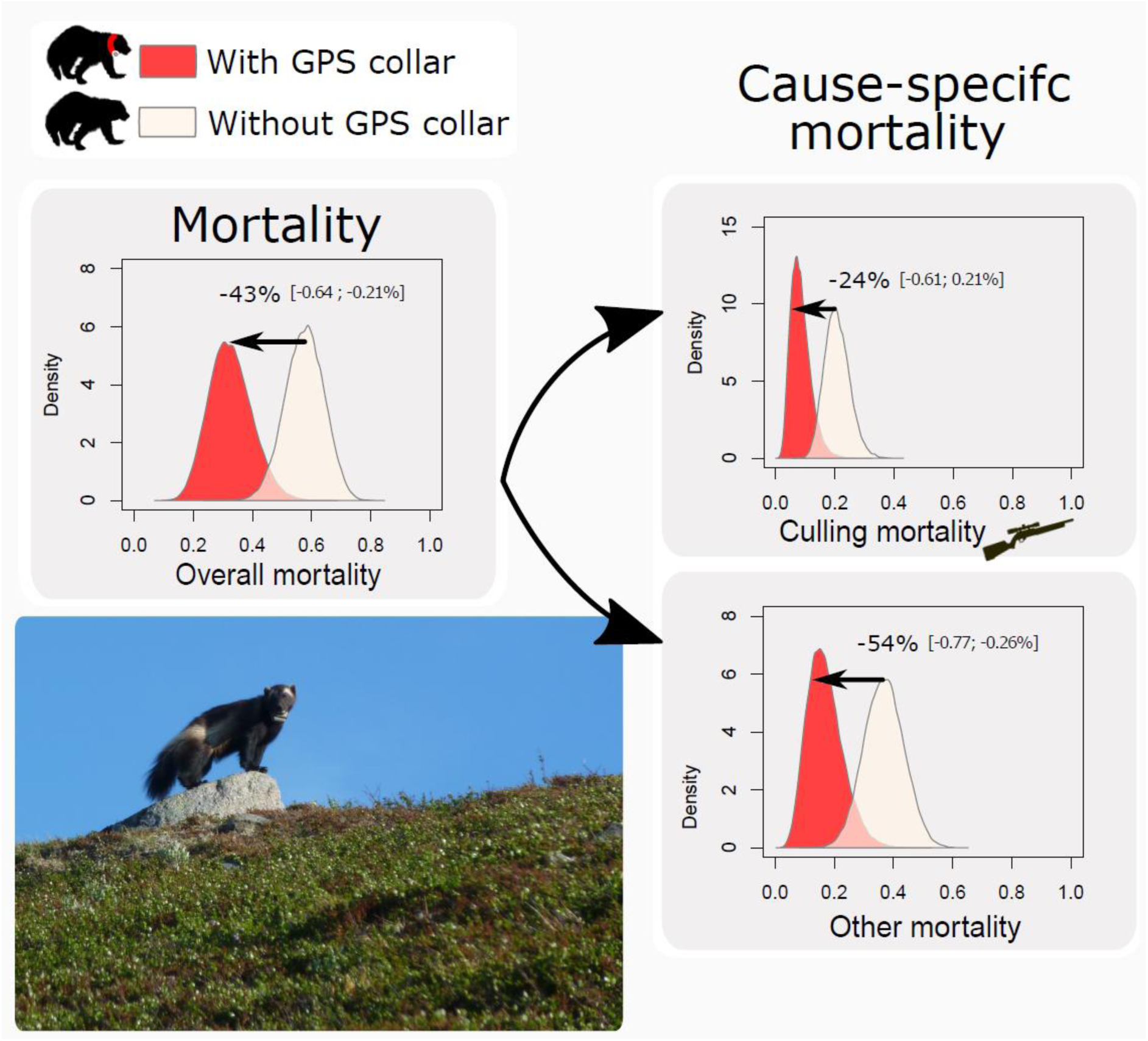
Posterior distributions of wolverine mortality rates with and without a GPS collar obtained using a Bayesian open population spatial capture-recapture model and data from non-invasive genetic sampling in Scandinavia. Proportional differences (95% credible interval) in mortality rates between individuals with and without a GPS collar are depicted for males in the northern area and between the monitoring season 2010/11-2011/12. Proportional differences in mortality rates between individuals with and without a GPS collar depended on the baseline probabilities (*h*0, *w*0) that varied with year, sex, and regions because the effect of capture on mortality rates (*βw*_*GPS*_, *βh*_*GPS*_) were quantified on the logit scale. The picture shows one of the studied wolverines wearing a GPS collar (©Thomas Strømseth).

The northern study area is situated in Finnmark and Troms County (70°10′ N, 24°70′) and the Central area in Trøndelag County (64°30′ N, 12°50′ E). Troms and Finnmark have a coastal alpine climate, while Trøndelag has a more continental climate. The northern area is dominated by alpine tundra and mountain birch forest (*Betula pubescens*) forms the tree line. Northern boreal forest, dominated by conifer (*Pinus sylvestris, Picea abies*), is common at lower elevations in the central area. High elevations of both areas are usually snow-covered from November to June. Free-ranging semi-domestic reindeer (*Rangifer tarandus*) is the most abundant ungulate in both areas. All reindeer herds migrate seasonally, but different husbandry practices and land use restrictions create variation in the timing, magnitude, and direction of the migrations. The coastal parts of Troms and Finnmark are almost completely devoid of reindeer in winter (Mattisson et al. 2011). In Trøndelag, seasonal migration is short, and most wolverine home ranges include both summer and winter reindeer-grazing areas. Free-ranging domestic sheep (*Ovis aries*) are present to a varying degree in both areas during summer.

### Non-invasive genetic sampling

Wolverines are monitored annually by Swedish and Norwegian authorities using non-invasive genetic sampling (NGS) and dead recoveries (Gervasi et al. 2016). Scat and hair samples were collected on snow along wolverine tracks from December to June, covering the entire distribution range of the species in Scandinavia. Non-invasive genetic samples were collected from individuals (≥1 year old), but not from young of the year. For further details on the DNA analysis procedure see (Flagstad et al. 2004, Brøseth et al. 2010, Gervasi et al. 2016). To ensure that the studied individuals with and without a GPS collars were exposed to similar conditions, we only considered non-invasive genetic samples collected within 70km of any genetic samples from collared individuals.

We used data from eight consecutive monitoring seasons (December-June) between 2009/10 (i.e. the winter when wolverine captures started) and 2016/17. After the winter 2016/17, none of the wolverines were wearing a GPS collar. The non-invasive dataset contained a total of 4989 (♀2446; ♂2543) genetically identified samples of which 2753 (♀1360; ♂1393) and 2236 (♀1183; ♂1053) were collected in the Central- and Northern areas, respectively (Appendix S1 Table S1). This corresponded to 1036 (♀555; ♂481) different individuals of which 616 (♀321; ♂295) and 421 (♀235; ♂186) were detected in the Central- and Northern areas, respectively (Appendix S1 Table S1). Note that only one individual was detected in both study areas. We also obtained recovery locations and genetic identification data from 424 (♀219; ♂ 205) individuals legally culled during the study period. Occasionally, individuals recovered dead due to other reasons (e.g. poaching, car collision) were also reported (♀6; ♂ 5).

### GPS covariate

All collared individuals were genetically identified from tissue collected during the capture which allowed us to match them with the NGS dataset. We created an individual and time-dependent binary covariate (*GPS*) coding 1 if the individual was wearing a GPS collar at any time during the monitoring season, and 0 otherwise. The latter included individuals that were never captured, and individuals before they were collared or after losing their collar. With the use of this covariate, we intended to account for any contemporary effect caused by the GPS collar.

### OPSCR model

To estimate the survival rates of wolverines, we built an open population spatial capture-recapture (OPSCR) model (Ergon and Gardner 2014, Bischof et al. 2016, Chandler et al. 2018), which addresses three challenges associated with the analyses of population-level NGS data: 1) detection is imperfect and heterogeneous in space and time (Kery and Schaub 2011), 2) individuals that reside primarily outside the surveyed area may be detected within it (Efford 2004), and 3) estimation of true survival rate (Ergon and Gardner 2014).

We estimated separate model parameters for each region and sex, except for the parameters that concerned the quantification of the differences between GPS-collared versus not GPS-collared individuals. Due to sample size limitations, we did not break estimates down further and assumed that differences between collared versus non-collared animals were consistent across regions and sex.

The OPSCR model is a hierarchical state-space model composed of three sub-models: 1) a model for detections during DNA searches, 2) a model for population dynamics, and 3). a model for density and movements.

#### The detection model

the detection model describes the relationship between individual detection probability and distance to its activity center (AC). We used the half-normal detection model that assumes that the probability *p* of detecting individual *i* at detector *j* and time *t* decreases with distance (D_*ijt*_) between the detector and the AC:

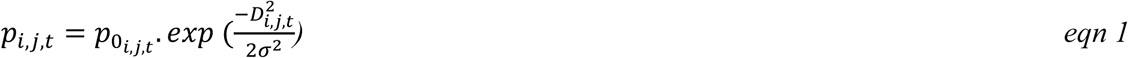

where p_0_ represents the detection probability at the location of the AC, and σ represents the width of the utilization distribution.

We placed a detector grid within a buffer of 70 km surrounding all detections of captured individuals for each region (Appendix S1, Figure 1). The spatial domain was defined as the detector grid surrounded by a 60 km buffer. We used grid cells as detectors and aggregated detections to the closest grid cell center. We used a partially aggregated binomial model that divided detectors (10 km resolution) into 25 subdetectors (2 km) and modeled the frequency of subdetectors with more than one detection as a binomial response with a sample size of 25 (for additional details see Milleret et al. 2018). To account for individual, spatial, and temporal heterogeneity in detection probability we included several linear effects on a logistic scale on the baseline detection probability (p_0_) (Appendix S2):

#### The multistate model

We used a multistate formulation for the population dynamics model (Lebreton and Pradel 2002) where each individual life history is represented by a succession of discrete states z_it_ that arises from a Markov process. Between two consecutive years, an individual *i* can either remain in its current state or transition to another one, with transition probabilities corresponding to vital rates. We considered four different states: “unborn” if the individual has not yet been recruited in the population. “alive” if it is alive. “dead legal” if it has died from legal culling between the start of the previous and current monitoring seasons. “Dead”: if it has a) died from any other cause of mortality between the start of the previous and current monitoring seasons or b) died earlier, regardless of the cause.

During the first occasion, individuals can only be designated as “unborn” or “alive” so that z_i,1_∼dcat(1-ψ, ψ, 0, 0), where ψ represents the probability to be already part of the population at t=1.

For *t≥2*, z_i,t_ is conditional on the state of individual *i* at *t-1:*

- If z_i,t-1_ =1, individual *i* is available to be recruited (transition to state 2), so z_i,t_ ∼dcat(1-γ_*t*_, γ_*t*_, 0, 0).
- If z_i,t-1_ =2 individual *i* can survive and remain z_it_=2 with probability Φ_t_, die from culling and transition to z_it_=3 with probability *h*_t_ or die from other causes and transition to z_i,t_ =4 with probability *h*_t_, so that z_i,t_ ∼ dcat(0, Φ_t_, h_t_, w_t_), where Φ_t_ = 1−h_t_ −w_t_
- Finally, all individuals in dead states (z_i,t-1_ = 3 or 4) transition to z_i,t-1_ = 4, the absorbing state, with probability 1.

To quantify differences in mortality rates between collared and non-collared individuals, we included the effect of the *GPS* covariate as a linear effect on the logistic scale for both culling (*h*) and other (*w*) mortality rates:

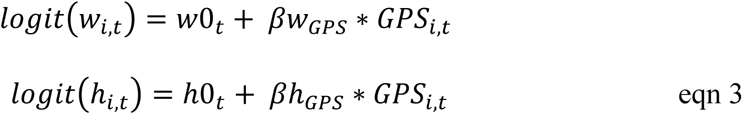

Note that we estimated separate *β* parameters for each region and sex.

The detection data y_ijt_ was modelled as the realization of a Binomial process based on the individual, detector, and time specific detection probability p_*i,j,t*_, the number of trials defined as the number subdetectors K, and conditional on individual state *z*_*i,t*_ (only individuals with the state “*alive*” can be detected.):

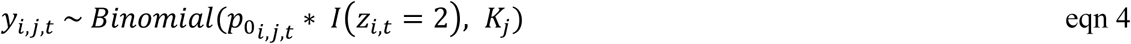

#### The movement model

We used an inhomogeneous binomial point process to model the distribution of individual ACs (Illian et al. 2008). This formulation allows the density of ACs to vary according to a spatial intensity function. ACs (*s*) at *t* = 1 were placed according to the intensity function λ(*s*_1_), that was based on a spatial covariate (*Dens;* locations of known dens from den visits conducted by authorities in their effort to document reproductions per 20 km habitat cell) that roughly reflects the species distributions 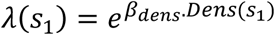. For *t≥2*, the intensity function was a combination of the intensity surface representing AC distribution (*β*_*Dens*_) and an isotropic multivariate normal distribution centered around the source coordinates (location of the AC at *t-1*) with standard deviation τ (see (Milleret et al. 2020) for more details). As a consequence, individual movement from year to year was modelled as a Markovian spatial point process. This feature of the model improves survival estimates as it takes into account the impact of animals moving within and out of the sampled area (Ergon and Gardner 2014, Gardner et al. 2018).

### Parameter estimation

We fitted the Bayesian OPSCR models using Markov chain Monte Carlo (MCMC) simulation with NIMBLE (de Valpine et al. 2017, NIMBLE Development Team 2019) in R version 3.3.3 (R Core Team 2018). We implemented different modelling techniques to increase MCMC efficiency (available in the nimbleSCR package (Bischof et al. 2020); see (Milleret et al. 2018, 2019, 2020, Turek et al. 2020) for details). We ran 4 chains, each with 42,500 iterations including a 12,500-iteration burn-in period. We considered models as converged when the Gelman-Rubin diagnostic (Rhat, (Gelman and Rubin 1992)) was < 1.1 for all parameters and by visually inspecting the trace plots. We summarized posterior distributions using their median and 95% credible interval (CrI).

## Results

Collared wolverines had a 47% (95% CrI: [21%;68%]) lower overall mortality rate compared to non-collared individuals (Figure 1). This difference was attributable mainly to lower mortality rate due to causes other than legal culling (*βw*_*GPS*_= −1.08 [−1.86; −0.46]), and to a lesser extent to a lower mortality rate due to culling (*βh*_*GPS*_= −0.37 [-1.09; 0.25]; Figure 1). On average, individuals with a collar had 58% [29; 80%] reduced probability to die from mortality due to causes other than legal culling than individuals without a GPS collar. Individuals wearing a GPS collar had a lower detectability rate compared to individuals without GPS collar 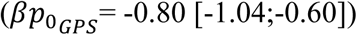. Additional details are presented in Appendix S1 Figure S2-6.

## Discussion

Using a single individual-based dataset containing wolverines both with and without a GPS collar, and an open population spatial capture-recapture model, we were able to provide a quantitative answer to a question that has eluded wildlife ecologists: can we extrapolate results obtained from a limited number of instrumented individuals to the population? We found that wolverines with a GPS collar had a lower mortality rate compared to individuals without a collar. Although collared individuals seemed to have slightly lower mortality associated with legal culling, the difference was mainly manifested through a lower mortality rate due to other causes. Our results call for prudence when extrapolating vital rate estimates from a sample of instrumented large carnivores to the population.

There are three main potential, not mutually exclusive, explanations for the observed difference in mortality between wolverines with and without GPS collar: a) wolverines selected for GPS-collaring had mortality that differed from the population average (“non-random sample”), b) instrumentation altered wolverine survival (“observer effect”), and c) that our NGS and OPSCR-based approach to mortality estimation inadvertently introduced a bias (“analytical artifact”). Being observational in nature, our study does not allow us to isolate an explanation unequivocally. However, we argue that the observer effect could be the primary cause for the observed difference in mortality between instrumented and non-instrumented animals.

### Analytical artifact

Our OPSCR model aimed at quantifying population dynamics of wolverines while accounting for imperfect detection inherent to NGS data. To reduce the chances of introducing a bias with our OPSCR model by inadvertently comparing different parts of the population, we restricted the spatial (Appendix S1, Figure S1) and temporal extents of the analysis to limit the differences in ecological conditions that individuals with and without collars are exposed to. Furthermore, we accounted for individual, temporal, and spatial heterogeneity in detectability in the population-level OPSCR model, including potential differences in detectability between GPS collared and non-collared individuals. All OPSCR model parameters, except parameters linked with the effect of wearing a GPS collar, were also sex- and region-specific thereby accounting for possibly important sources of variation.

### Non-random sampling

Non-invasive genetic sampling and the collaring of wolverines targeted all segments of the population (except cubs of the year). Captures from helicopter also occurred opportunistically by following any fresh wolverine tracks encountered on snow, without intentional focus on a specific part of the population. Captures may have been unintentionally biased towards individuals whose tracks are likely to be encountered (e.g., animals moving in open areas and traveling longer distances) and those that are more vulnerable to capture once detected (e.g. young individuals). Indeed, the proportion of 1-year old among collared individuals (41%) was higher than their expected prevalence in the population (29%; Landa et al. 1998). With a negative age-mortality relationship (Persson et al. 2009)(Persson et al. 2009), the proportion of young individuals among GPS-collared individual being higher than expected for this wolverine population, we would expect higher mortality among GPS tagged animals than the population average. Yet, we detected the opposite effect, which suggests that the explanation for the mortality difference lies elsewhere.

### Observer effect

A third possibility is that wearing a GPS collar itself impacts mortality of instrumented wolverines. Mortality due to other causes includes three main causes that our study can not differentiate: 1) natural (e.g., age, diseases, intra and inter-specific killing), 2) traffic, and 3) illegal killing. The literature generally reports negative or neutral effect of the capture, handling, and wearing of bio-logger on the survival of individuals (Sergio et al. 2015, Bodey et al. 2018), and we are not aware of any mechanisms that could give a competitive advantage to GPS collared wolverines to a magnitude that would decrease their natural or traffic mortality rates. Although patterns of carnivore mortality due to causes other than culling remain cryptic, we know that illegal culling account for a large part of carnivore mortality in Scandinavia (Persson et al. 2009, Liberg et al. 2011, Milleret et al. 2016, Rauset et al. 2016). Therefore, a plausible explanation for the observed result is that GPS collars could shield individuals against illegal killing. First, the collar in itself can act as a deterrent as the likelihood of detecting poaching events increases. Secondly, the wolverines were captured as part of a wildlife-human conflict project studying predation on semi-domestic reindeer and sheep (e.g. Mattisson et al. 2016). In Norway, compensation for livestock losses to carnivores is estimated based on the number of detected domestic preys that could be documented as killed by a carnivore. As information provided by the GPS collar is partly used to conduct predation studies, it generally results in a higher number of documented kills, which could give an incentive to keep GPS collared individuals alive.

Instrumented versus non-instrumented individuals have been suggested to have different survival rates in other large carnivore populations (Schmidt et al. 2015, Borg et al. 2016, Treves et al. 2017), but evaluating the representativeness of instrumented individuals is particularly challenging as it also requires to obtain data on non-instrumented individuals. Using a comprehensive spatially explicit population-level model taking into account imperfect detection inherent to non-invasive genetic sampling, we showed that GPS collared large carnivores may not always constitute a representative sample of the population for the estimation of survival rates. Although our OPSCR model would greatly benefit from incorporating age structure, age information was not available for individuals detected solely with NGS. To our knowledge, this study is the first to compare survival rates of carnivores with and without telemetry collars, using an independent source of data collected at the individual level, and a comprehensive statistical framework that takes into account imperfect detection. Regardless of the exact mechanisms behind the non-representativeness of the instrumented sample, it has important implication for management and conservation, as many studies use data from GPS collared individuals to draw population-level inferences (Hebblewhite and Haydon 2010) and to parameterize population level models (Liberg et al. 2011)..

## Authors contribution

C.M developed the concept of this study with input from R.B and P.D. J.M and J.O provided data on captured wolverines. Wolverine genetic data extraction and preparation were coordinated by H.B. C.M led the analysis with help from R.B and P.D. C.M led the writing with contributions from R.B, P.D, H.B, J.O and J.M. All authors gave final approval for publication.

## Data

Data, R script and nimble codes are available on github repository https://github.com/Cyril-Milleret/Public/tree/master/WolverineGPSSurvival

## Acknowledgment

This work was funded by the Norwegian Environment Agency (Miljødirektoratet), the Swedish Environmental Protection Agency (Naturvårdsverket), and the Research Council of Norway (NFR 286886). The County administrations in Trøndelag, Troms, and Finnmark, the Carnivore Management Boards in regions 6 and 8, and the Reindeer Development Fund in Norway also provided financial assistance. We thank the field staff from the State Nature Inspectorate and members of the public that collected the monitoring data for the large carnivore database Rovbase3.0 (rovbase.no, rovbase.se). Computations/simulations were performed on resources provided by NMBU’s computing cluster “Orion”. A special thanks to P. Segerström, E. Segerström, T. Strømseth, and J. M. Arnemo for catching and collaring the wolverines, and the State Nature Inspectorate, reindeer herders, and all additional people that have contributed to captures and data collection in the field. We also thanks P. de Valpine and D. Turek for their assistance with NIMBLE.

## Appendix S1

**Table S1.** Summary of the number of non-invasive genetic samples from wolverines and individuals genetically identified from 2009/10-2016/17 in the Central and Northern study areas (see Figure 1) for females (F) and males (M).

| Year | 2009 | 2010 | 2011 | 2012 | 2013 | 2014 | 2015 | 2016 | Total |
| --- | --- | --- | --- | --- | --- | --- | --- | --- | --- |
| **N detections** |  |  |  |  |  |  |  |  |  |
| **Sex** | **Central** |  |  |  |  |  |  |  |  |
| **M** | 129 | 109 | 145 | 176 | 177 | 187 | 213 | 257 | 1393 |
| **F** | 169 | 108 | 149 | 149 | 199 | 162 | 212 | 212 | 1360 |
| **North** |  |  |  |  |  |  |  |  |  |
| **M** | 124 | 113 | 148 | 116 | 168 | 98 | 121 | 165 | 1053 |
| **F** | 124 | 99 | 167 | 107 | 182 | 139 | 134 | 231 | 1183 |
| **N individuals detected** |  |  |  |  |  |  |  |  |  |
| **Central** |  |  |  |  |  |  |  |  |  |
| **M** | 55 | 49 | 64 | 70 | 61 | 64 | 72 | 73 | 508 |
| **F** | 71 | 53 | 67 | 75 | 80 | 73 | 87 | 91 | 597 |
| **North** |  |  |  |  |  |  |  |  |  |
| **M** | 50 | 39 | 49 | 34 | 43 | 32 | 31 | 35 | 313 |
| **F** | 55 | 48 | 77 | 48 | 69 | 62 | 59 | 72 | 490 |

**Figure S1.**
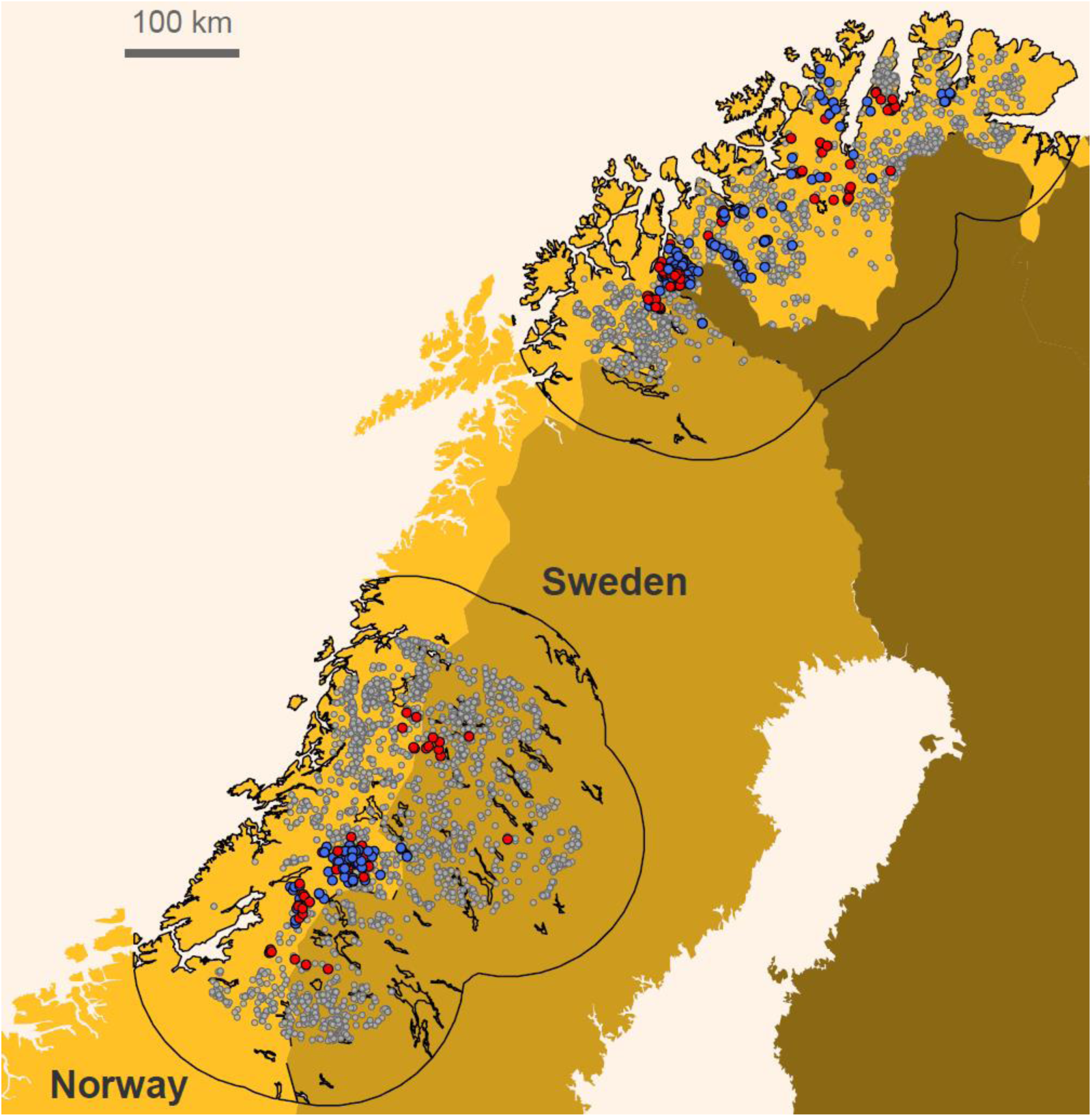
Locations of non-invasive genetic samples of wolverines collected during eight consecutive monitoring seasons (December– June; 2009/10-2016/17) in Sweden and Norway. Blue and red dots show genetically identified samples from male and female individuals with GPS-collars, respectively. Grey dots represent detections from all other individuals. Polygons with a black outline represent the two study areas (North and Center) and include the buffer (area of the polygon without detections) considered in the open-population spatial capture-recapture analysis.

**Figure S2.**
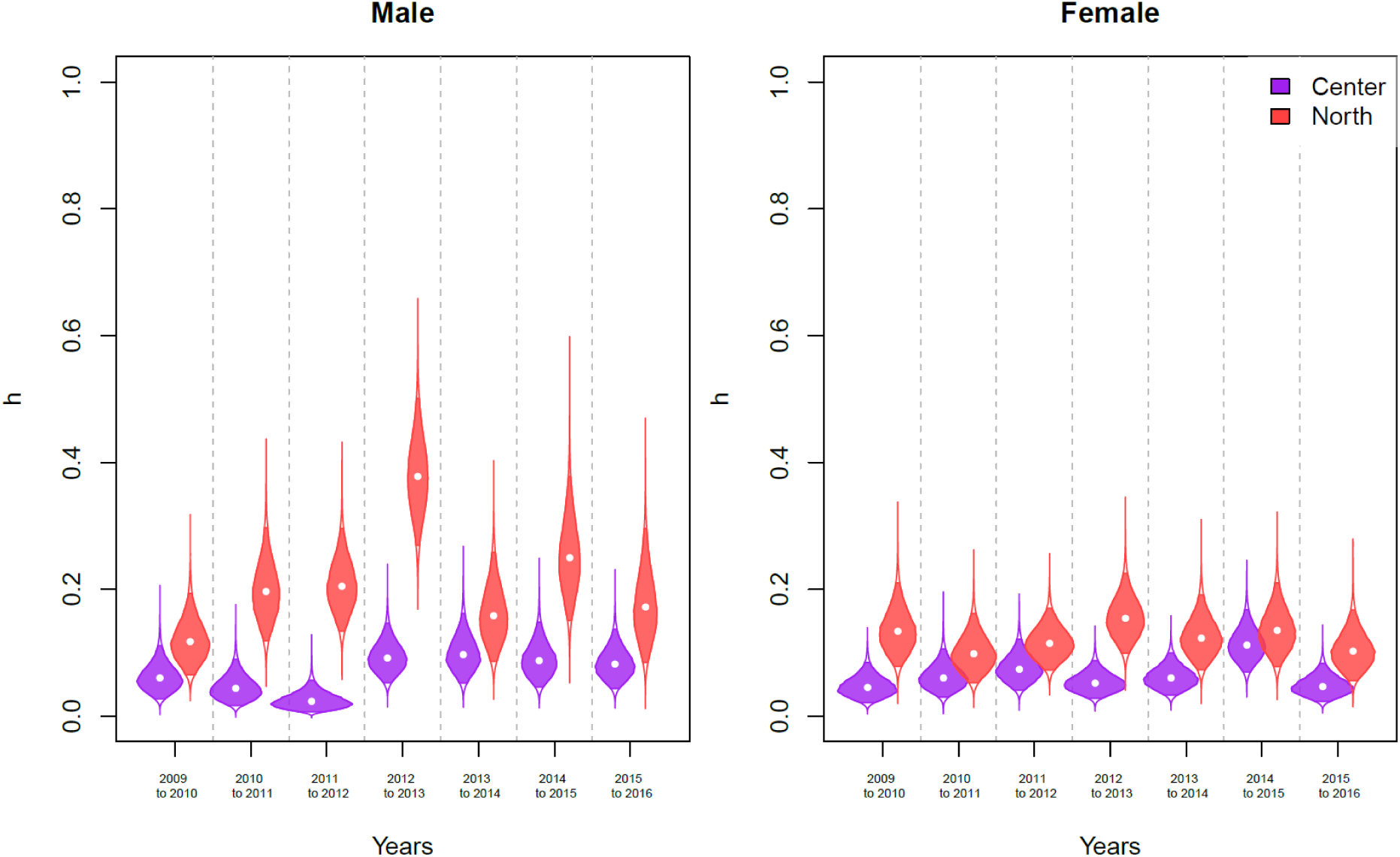
Mortality probabilities due to legal culling (*h*) for male and female wolverines. Shown are overall estimates throughout the Central and Northern study areas (Figure 1). Violins show the posterior distribution of mortality estimates (points: median estimates; solid colors: 95% credible interval). Estimates reflect mortalities occurring between the start of one sampling season and the start of the next.

**Figure S3.**
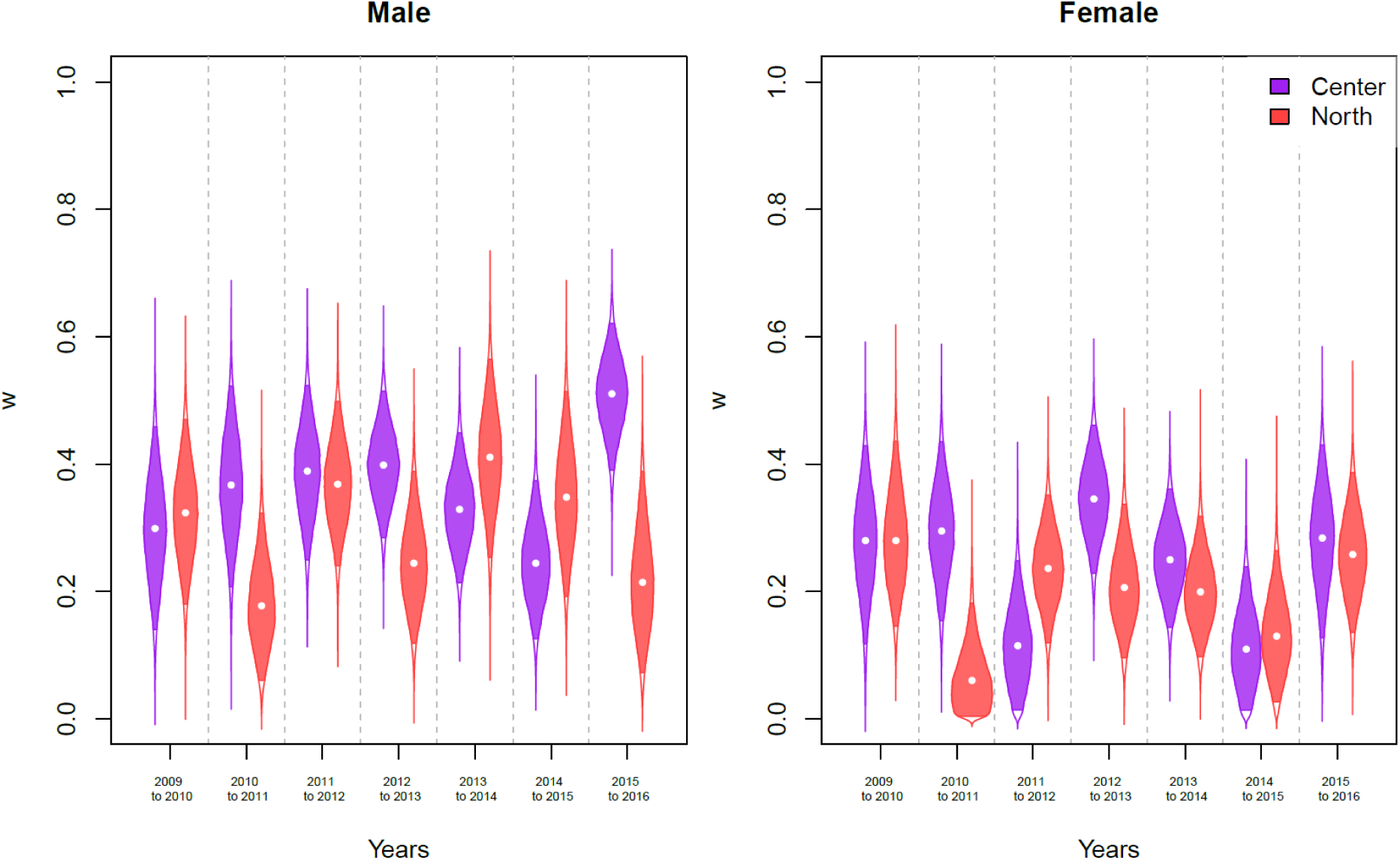
Mortality probabilities due to causes other than legal culling (*w*) for male and female wolverines. Shown are overall estimates throughout the Central and Northern study areas (Figure 1). Violins show the posterior distribution of mortality estimates (points: median estimates; solid colors: 95% credible interval). Estimates reflect mortalities occurring between the start of one sampling season and the start of the next.

**Figure S4.**
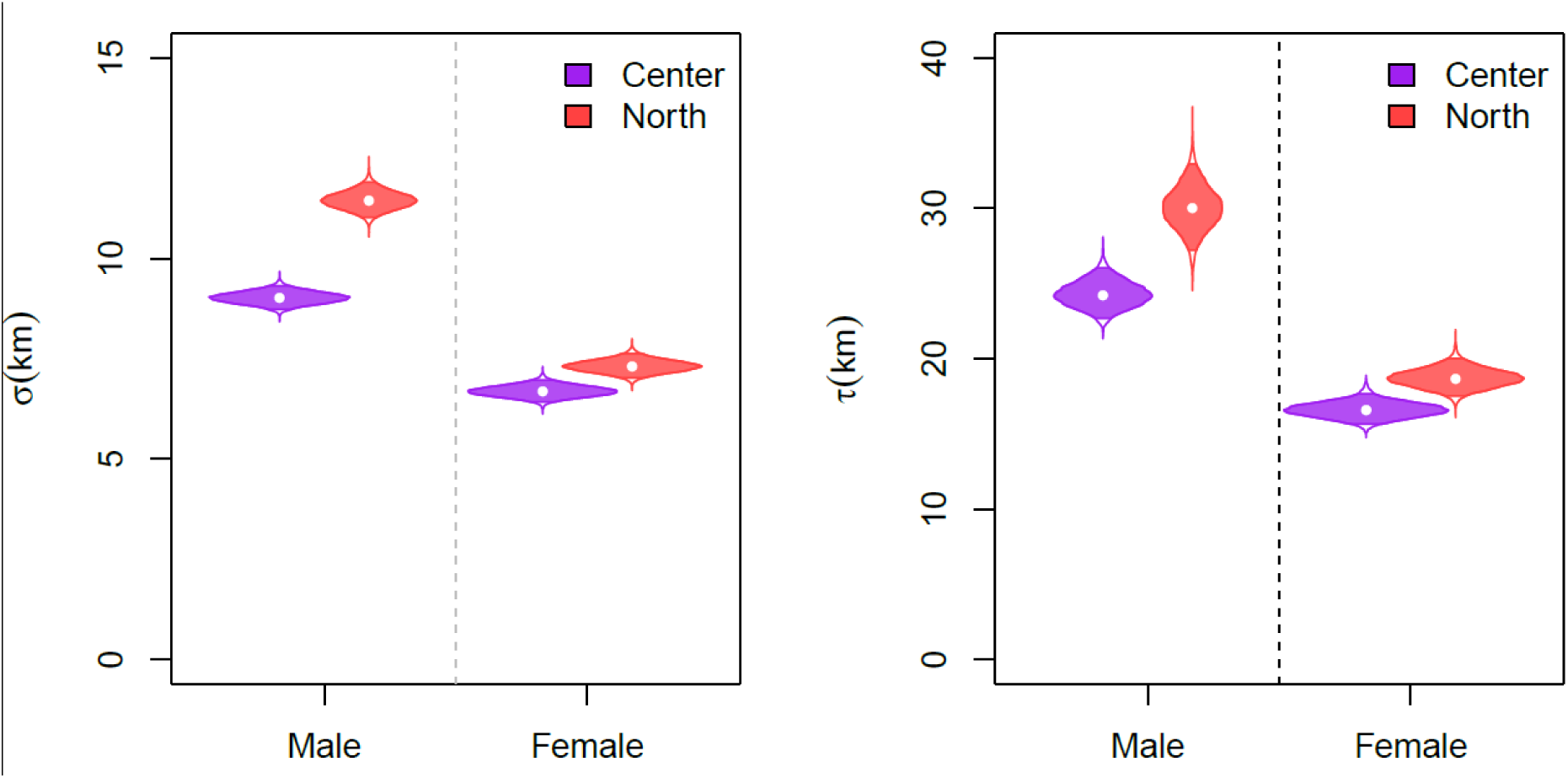
Estimates of the scale parameter *σ* of the detection function representing range of movement of individuals around their activity centers (AC), and scale parameter τ of the bivariate normal representing inter-annual movement of AC locations. Estimates are presented for male and female wolverines for the Central and Northern study areas (Figure 1). Violins show the posterior distribution estimates (points: median estimates; solid colors: 95% credible interval).

**Figure S5.**
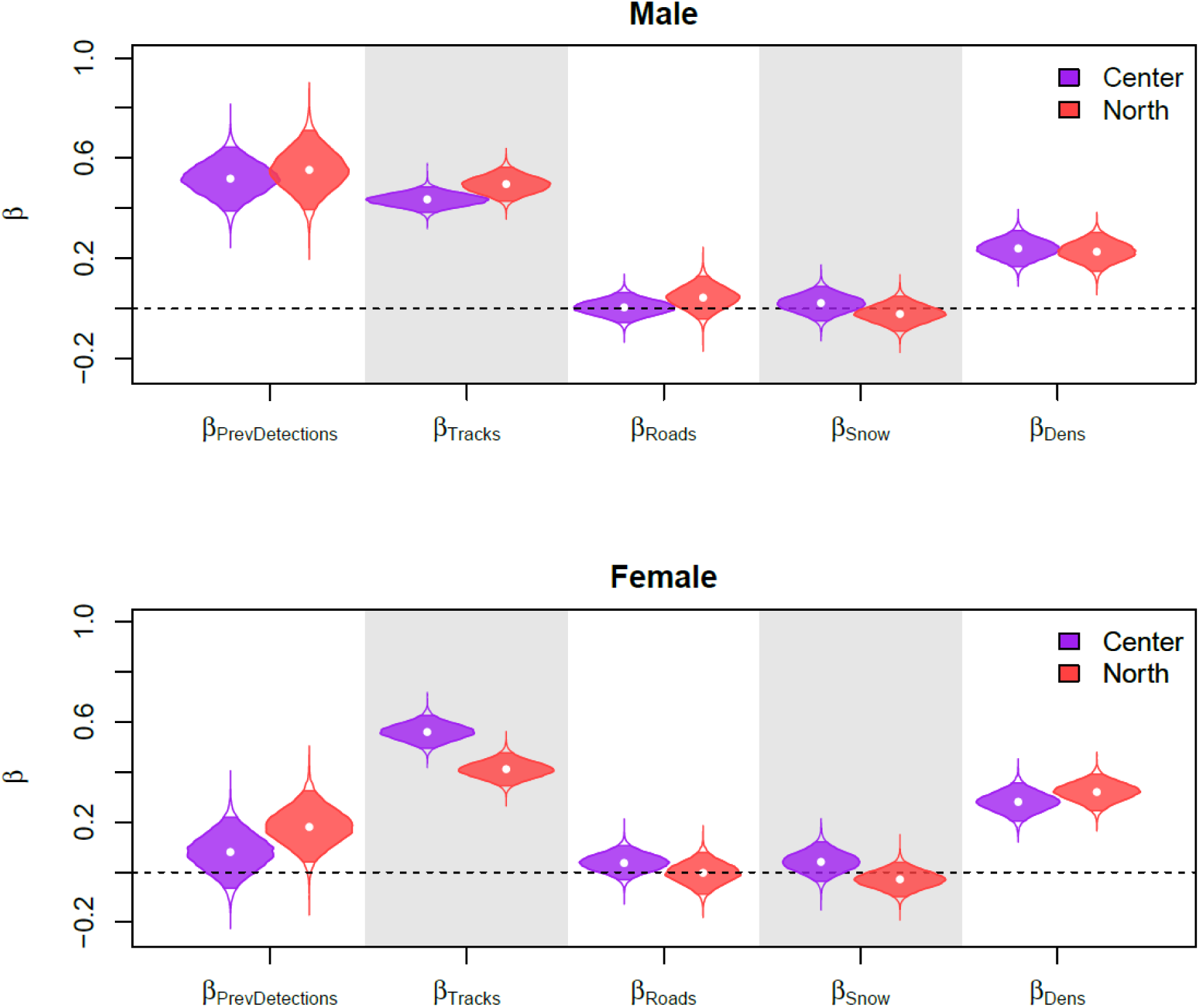
Beta coefficients to model the logit linear effect of spatial covariates on the baseline detection probability (p_0;_ *β*_*PrevDetections*_, *β*_*Tracks*_, *β*_*Roads*_, *β*_*Snow*_; see OPSCR model section for further details) and density (*β*_*Dens*_). Estimates are presented for male and female wolverines for the Central and Northern study areas (Figure 1). Violins show the posterior distribution estimates (points: median estimates; solid colors: 95% credible interval).

**Figure S6.**
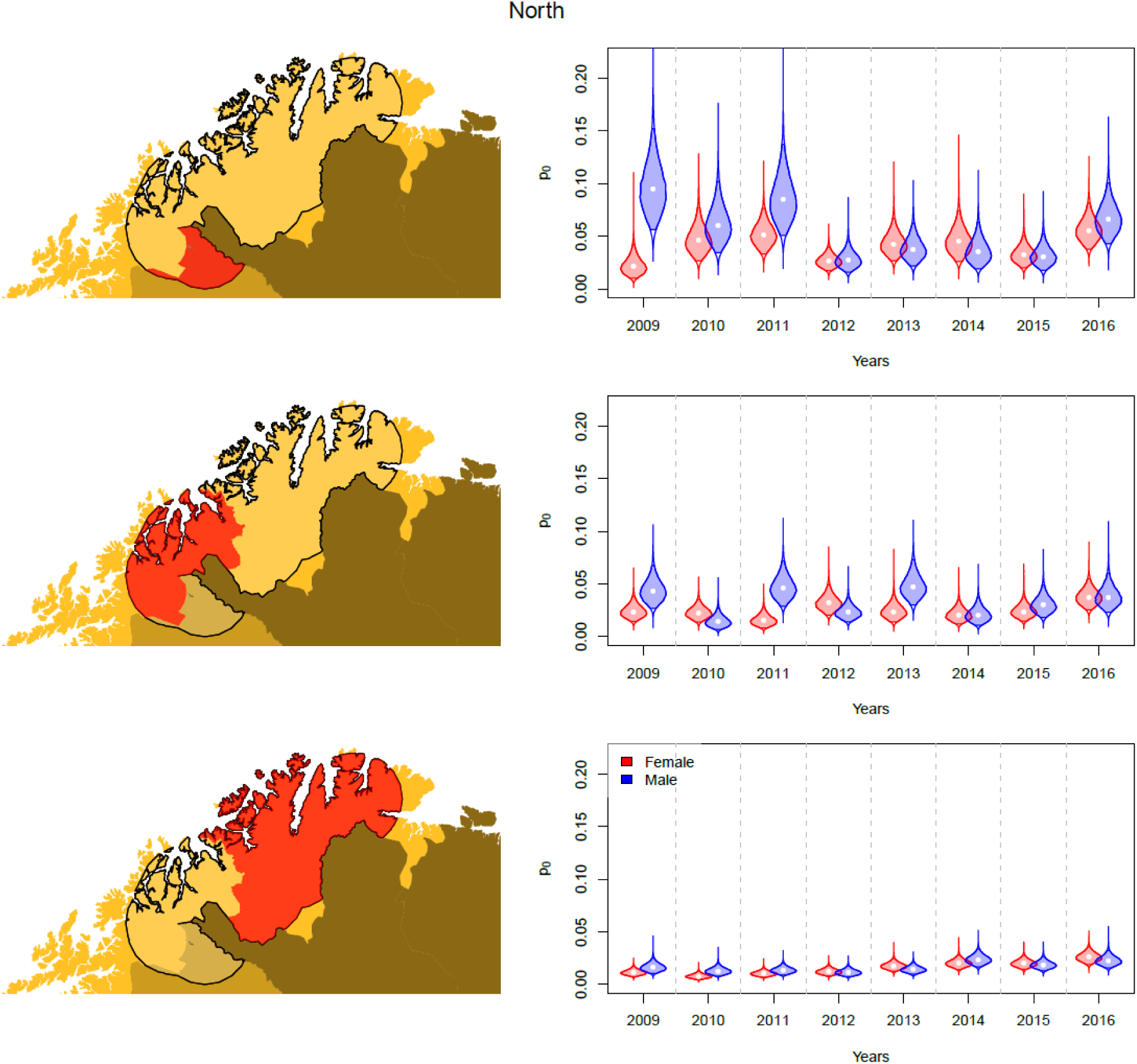
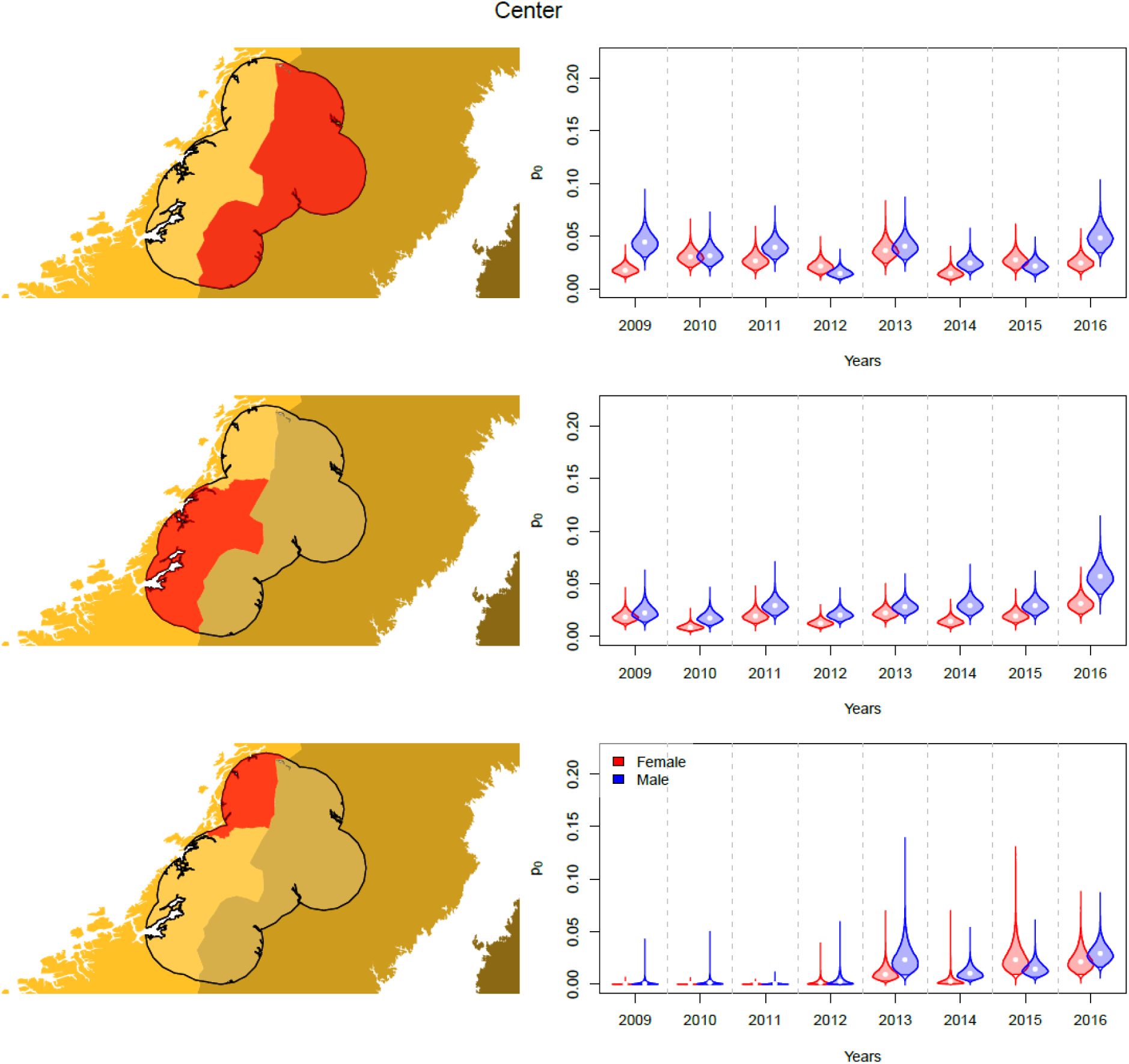
Estimates of the baseline detection probability (*p*_*0*_) presented for male and female wolverines for the central and northern study areas (Figure 1). *p*_*0*_ was estimated separately for each year and for every region (three for the Central and Northern study areas, colored red in the map from the left panel plot). Violins show the posterior distribution estimates (points: median estimates; solid colors: 95% credible interval).

## Appendix S2

To account for individual, spatial, and temporal heterogeneity in detection probability we included several linear effects on a logistic scale on the baseline detection probability (p_0_):

1. Length of GPS search tracks logged by searchers within each detector grid cell in each monitoring period (*Tracks*).
2. Average distance from the nearest road (*Roads*): the distance from each detector to the closest road (1:100,000, the Swedish mapping, cadastral and land registration authority; N50 kartdata, the Norwegian Mapping Authority,). This variable represents accessibility, which we predicted to facilitate detectability.
3. Yearly average percentage of snow cover in each detector grid cell (MODIS at 0.1 degrees resolution, www.neo.sci.gsfc.nasa.gov, accessed 2019-10-11) between December 1– June 31 (*Snow*). As wolverine NGS during winter relies heavily on the presence of snow, we predicted that greater snow cover increases detectability.
4. Indicator of whether an individual was detected or not during the previous monitoring season (*PrevDetection*). Previous detection could be expected to positively influence the probability of being detected at subsequent occasions (Gervasi et al. 2014).
5. Whether the individual was GPS collared (*GPS*) or not to account for different detection probabilities. Furthermore, we estimated independent baseline detection probabilities (*p*_0*Intercept*_) for each year (*t*):

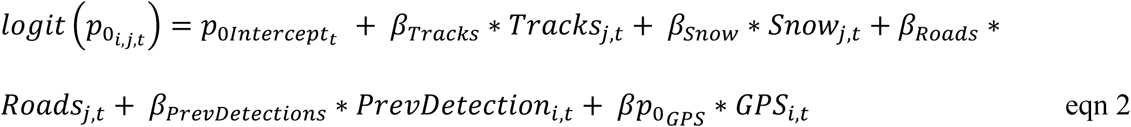

Note that we estimated separate *β* parameters for each region and sex.

## References

Arnemo, J. M., A. L. Evans, and A. Fahlman. 2012. Biomedical protocols for free-ranging brown bears, gray wolves, wolverines and lynx. Hedmark University College. Evenstad, Norway.

Bischof, R., H. Brøseth, and O. Gimenez. 2016. Wildlife in a Politically Divided World: Insularism Inflates Estimates of Brown Bear Abundance. Conservation Letters 9:122–130.

Bischof, R., J. G. O. Gjevestad, A. Ordiz, K. Eldegard, and C. Milleret. 2019. High frequency GPS bursts and path-level analysis reveal linear feature tracking by red foxes. Scientific reports 9:1–13.

Bischof, R., D. Turek, C. Milleret, T. Ergon, P. Dupont, and P. de Valpine. 2020. nimbleSCR: Spatial Capture-Recapture (SCR) Methods Using “nimble.”

Bodey, T. W., I. R. Cleasby, F. Bell, N. Parr, A. Schultz, S. C. Votier, and S. Bearhop. 2018. A phylogenetically controlled meta-analysis of biologging device effects on birds: Deleterious effects and a call for more standardized reporting of study data. Methods in Ecology and Evolution 9:946–955.

Borg, B. L., S. M. Arthur, N. A. Bromen, K. A. Cassidy, R. McIntyre, D. W. Smith, and L. R. Prugh. 2016. Implications of harvest on the boundaries of protected areas for large carnivore viewing opportunities. PLoS one 11:e0153808.

Brøseth, H., Ø. Flagstad, C. Wärdig, M. Johansson, and H. Ellegren. 2010. Large-scale noninvasive genetic monitoring of wolverines using scats reveals density dependent adult survival. Biological Conservation 143:113–120.

Canine, N. G. 1990. Unrecognized anti-predator behaviour can bias observational data. Animal Behaviour 39:195–197.

Carricondo-Sanchez, D., B. Zimmermann, P. Wabakken, A. Eriksen, C. Milleret, A. Ordiz, A. Sanz-Pérez, and C. Wikenros. 2020. Wolves at the door? Factors influencing the individual behavior of wolves in relation to anthropogenic features. Biological Conservation 244:108514.

Cattet, M., J. Boulanger, G. Stenhouse, R. A. Powell, and M. J. Reynolds-Hogland. 2008. An evaluation of long-term capture effects in ursids: implications for wildlife welfare and research. Journal of Mammalogy 89:973–990.

Chandler, R. B., J. Hepinstall-Cymerman, S. Merker, H. Abernathy-Conners, and R. J. Cooper. 2018. Characterizing spatio-temporal variation in survival and recruitment with integrated population models. The Auk 135:409–426.

Cooke, S. J., V. M. Nguyen, S. T. Kessel, N. E. Hussey, N. Young, and A. T. Ford. 2017. Troubling issues at the frontier of animal tracking for conservation and management. Conservation Biology 31:1205–1207.

Darrow, P. A., and J. A. Shivik. 2009. Bold, shy, and persistent: Variable coyote response to light and sound stimuli. Applied Animal Behaviour Science 116:82–87.

Efford, M. 2004. Density estimation in live-trapping studies. Oikos 106:598–610.

Ergon, T., and B. Gardner. 2014. Separating mortality and emigration: modelling space use, dispersal and survival with robust-design spatial capture-recapture data. Methods in Ecology and Evolution 5:1327–1336.

Esteruelas, N. F., N. Huber, A. L. Evans, A. Zedrosser, M. Cattet, F. Palomares, M. Angel, J. E. Swenson, and J. M. Arnemo. 2016. Leukocyte coping capacity as a tool to assess capture- and handling-induced stress in scandinavian brown bears (ursus arctos). Journal of Wildlife Diseases 52:S40–S53.

Flagstad, Ø., E. V. A. Hedmark, A. Landa, H. Brøseth, J. Persson, R. Andersen, P. Segerström, and H. Ellegren. 2004. Colonization history and noninvasive monitoring of a reestablished wolverine population. Conservation Biology 18:676–688.

Gardner, B., R. Sollmann, N. S. Kumar, D. Jathanna, and K. U. Karanth. 2018. State space and movement specification in open population spatial capture-recapture models. Ecology and Evolution 0.

Gelman, A., and D. B. Rubin. 1992. Inference from iterative simulation using multiple sequences. Statistical Science 7:457–511.

Gervasi, V., H. Brøseth, O. Gimenez, E. B. Nilsen, and J. D. C. Linnell. 2014. The risks of learning: confounding detection and demographic trend when using count-based indices for population monitoring. Ecology and Evolution 4:4637–4648.

Gervasi, V., H. Brøseth, O. Gimenez, E. B. Nilsen, J. Odden, Ø. Flagstad, and J. D. C. Linnell. 2016. Sharing data improves monitoring of trans-boundary populations: the case of wolverines in central Scandinavia. Wildlife Biology 22:95–106.

Hebblewhite, M., and D. T. Haydon. 2010. Distinguishing technology from biology: a critical review of the use of GPS telemetry data in ecology. Philosophical Transactions of the Royal Society B: Biological Sciences 365:2303–2312.

Illian, J., A. Penttinen, H. Stoyan, and D. Stoyan. 2008. Statistical analysis and modelling of spatial point patterns. John Wiley & Sons.

Kery, M., and M. Schaub. 2011. Bayesian Population Analysis using WinBUGS: A Hierarchical Perspective. Elsevier Science.

Landa, A., J. Tufto, R. Franzén, T. Bø, M. Lindén, and J. E. Swenson. 1998. Active wolverine Gulo gulo dens as a minimum population estimator in Scandinavia. Wildlife Biology 4:159–168.

Lebreton, J. D., and R. Pradel. 2002. Multistate recapture models: Modelling incomplete individual histories. Journal of Applied Statistics 29:353–369.

Lechenne, M., J. Arnemo, C. Brøjer, H. Andrén, and E. Agren. 2012. Mortalities due to constipation and dystocia caused by intraperitoneal radio-transmitters in Eurasian lynx (Lynx lynx). European Journal of Wildlife Research 58.

Liberg, O., Å. Aronson, S. M. Brainerd, J. Karlsson, H.-C. Pedersen, H. Sand, and P. Wabakken. 2010. Integrating research into management of a recolonizing wolf population – the Scandinavian model. Page in M. Musiani, L. Boitani, and P. Paquet, editors. (Eds.). The World of Wolves: New perspectives on ecology, behaviour and policy. University of Calgary Press, Calgary, Alberta, Canada.

Liberg, O., G. Chapron, P. Wabakken, H. C. Pedersen, N. T. Hobbs, and H. Sand. 2011. Shoot, shovel and shut up: cryptic poaching slows restoration of a large carnivore in Europe. Proceedings of the Royal Society B: Biological Sciences 279:910–915.

MacFarlane, G., and S. King. 2002. Observer presence influences behaviour of the semaphore crab, Heloecious cordiformis. Animal Behaviour – ANIM BEHAV 63:1191–1194.

Mattisson, J., J. Odden, E. B. Nilsen, J. D. C. Linnell, J. Persson, and H. Andrén. 2011. Factors affecting Eurasian lynx kill rates on semi-domestic reindeer in northern Scandinavia: Can ecological research contribute to the development of a fair compensation system? Biological Conservation 144:3009–3017.

Mattisson, J., G. R. Rauset, J. Odden, H. Andrén, J. D. C. Linnell, and J. Persson. 2016. Predation or scavenging? Prey body condition influences decision-making in a facultative predator, the wolverine. Ecosphere 7:e01407.

Mech, L. D., and S. M. Barber. 2002. A critique of wildlife radio-tracking and its use in national parks. Northern Prairie Wildlife Research Center.

Meeuwig, J. J., R. G. Harcourt, and F. G. Whoriskey. 2015. When Science Places Threatened Species at Risk. Conservation Letters 8:151–152.

Milleret, C., P. Dupont, C. Bonenfant, H. Brøseth, Ø. Flagstad, C. Sutherland, and R. Bischof. 2019. A local evaluation of the individual state-space to scale up Bayesian spatial capture–recapture. Ecology and Evolution 9:352–363.

Milleret, C., P. Dupont, H. Brøseth, J. Kindberg, A. Royle J., and R. Bischof. 2018. Using partial aggregation in spatial capture recapture. Methods in Ecology and Evolution 0.

Milleret, C., P. Dupont, J. Chipperfield, D. Turek, H. Brøseth, O. Gimenez, P. de Valpine, and R. Bischof. 2020. Estimating abundance with interruptions in data collection using open population spatial capture-recapture models. Ecosphere 11:e03172.

Milleret, C., P. Wabakken, O. Liberg, M. Åkesson, Ø. Flagstad, H. P. Andreassen, and H. Sand. 2016. Let’s stay together? Intrinsic and extrinsic factors involved in pair bond dissolution in a recolonizing wolf population. Journal of Animal Ecology.

NIMBLE Development Team. 2019. NIMBLE: MCMC, Particle Filtering, and Programmable Hierarchical Modeling. https://cran.r-project.org/package=nimble.

Packer, C., S. Shivakumar, V. Athreya, M. E. Craft, H. Dhanwatey, P. Dhanwatey, B. Gurung, A. Joshi, H. Kushnir, J. D. C. Linnell, and N. M. Fountain-Jones. 2019. Species-specific spatiotemporal patterns of leopard, lion and tiger attacks on humans. Journal of Applied Ecology 56:585–593.

Persson, J., G. Ericsson, and P. Segerström. 2009. Human caused mortality in the endangered Scandinavian wolverine population. Biological conservation 142:325–331.

R Core Team. 2018. R: A language and environment for statistical computing. Vienna,. Austria: R Foundation for Statistical Computing.

Rauset, G. R., H. Andrén, J. E. Swenson, G. Samelius, P. Segerström, A. Zedrosser, and J. Persson. 2016. National Parks in Northern Sweden as Refuges for Illegal Killing of Large Carnivores. Conservation Letters.

Schmidt, J. H., D. S. Johnson, M. S. Lindberg, and L. G. Adams. 2015. Estimating demographic parameters using a combination of known-fate and open N-mixture models. Ecology 96:2583–2589.

Sergio, F., G. Tavecchia, A. Tanferna, L. López Jiménez, J. Blas, R. De Stephanis, T. A. Marchant, N. Kumar, and F. Hiraldo. 2015. No effect of satellite tagging on survival, recruitment, longevity, productivity and social dominance of a raptor, and the provisioning and condition of its offspring. Journal of Applied Ecology 52:1665–1675.

Treves, A., J. A. Langenberg, J. V López-Bao, and M. F. Rabenhorst. 2017. Gray wolf mortality patterns in Wisconsin from 1979 to 2012. Journal of mammalogy 98:17–32.

Turek, D., C. Milleret, T. Ergon, H. Brøseth, and P. de Valpine. 2020. Efficient Estimation of Large-Scale Spatial Capture-Recapture Models. bioRxiv.

de Valpine, P., D. Turek, C. J. Paciorek, C. Anderson-Bergman, D. T. Lang, and R. Bodik. 2017. Programming with models: writing statistical algorithms for general model structures with NIMBLE. Journal of Computational and Graphical Statistics 26:403–413.

Young, J. K., L. Touzot, and S. P. Brummer. 2019. Persistence and conspecific observations improve problem-solving abilities of coyotes. PloS one 14:e0218778.

